# CryoGAN: A New Reconstruction Paradigm for Single-Particle Cryo-EM *via* Deep Adversarial Learning

**DOI:** 10.1101/2020.03.20.001016

**Authors:** Harshit Gupta, Michael T. McCann, Laurène Donati, Michael Unser

**Author notes:** Equal contribution (Harshit Gupta, Michael T. McCann).

## Abstract

We present CryoGAN, a new paradigm for single-particle cryo-EM reconstruction based on unsupervised deep adversarial learning. The major challenge in single-particle cryo-EM is that the imaged particles have unknown poses. Current reconstruction techniques are based on a marginalized maximum-likelihood formulation that requires calculations over the set of all possible poses for each projection image, a computationally demanding procedure. CryoGAN sidesteps this problem by using a generative adversarial network (GAN) to learn the 3D structure that has simulated projections that most closely match the real data in a distributional sense. The architecture of CryoGAN resembles that of standard GAN, with the twist that the generator network is replaced by a model of the cryo-EM image acquisition process. CryoGAN is an unsupervised algorithm that only demands projection images and an estimate of the contrast transfer function parameters. No initial volume estimate or prior training is needed. Moreover, CryoGAN requires minimal user interaction and can provide reconstructions in a matter of hours on a high-end GPU. In addition, we provide sound mathematical guarantees on the recovery of the correct structure. CryoGAN currently achieves a 8.6 Å resolution on a realistic synthetic dataset. Preliminary results on real *β*-galactosidase data demonstrate CryoGAN’s ability to exploit data statistics under standard experimental imaging conditions. We believe that this paradigm opens the door to a family of novel likelihood-free algorithms for cryo-EM reconstruction.

---

Single-particle cryo-electron microscopy (cryo-EM) is a powerful method to determine the atomic structure of macro-molecules by imaging them with electron rays at cryogenic temperatures [1–3].

Its popularity has rocketed in recent years, culminating in 2017 with the Nobel Prizes of Jacques Dubochet, Richard Henderson, and Joachim Frank. There exists a multitude of software packages to produce high-resolution 3D structure(s) from 2D measurements [4–11]. These sophisticated algorithms, which include projection-matching approaches and maximum-likelihood optimization frameworks, enable the determination of structures with unprecedented atomic resolution.

Yet reconstruction procedures in single-particle cryo-EM still face complex obstacles. The task involves a high-dimensional, nonconvex optimization problem with numerous local minima. Hence, the outcome of the global process is predicated on the quality of the initial reconstruction [12, 13]. Moreover, one still often relies on the input of an expert user for appropriate processing decisions and parameter tuning [14]. Even for more automated methods, the risk of outputting incorrect and misleading 3D reconstructions is ever-present. A key reason behind such complexity is that the imaged particles have unknown poses. To handle this, most software packages rely on a marginalized maximum-likelihood (ML) formulation [15] that is solved through an expectation-maximization algorithm [9, 11]. The latter involves calculations over the discretized space of poses for each projection, a computationally demanding procedure.

To bypass these limitations, we introduce CryoGAN, an unsupervised reconstruction algorithm for single-particle cryo-EM that exploits the remarkable ability of generative adversarial networks (GANs) to capture data distributions [16]. Similar to GANs, CryoGAN is driven by the competitive training of two entities: one that tries to capture the distribution of real data, and another that discriminates between generated samples and samples from the real dataset (Figure 1). In a classical GAN, the two entities are each a convolutional neural network (CNN). They are known as the generator and the discriminator and are trained simultaneously using backpropagation. The important twist with CryoGAN is that we replace the generator network by a cryo-EM physics simulator. By doing so, CryoGAN learns the 3D density map whose simulated projections are the most consistent with a given dataset of 2D measurements in a distributional sense (see *Online Methods - Mathematical Framework*).

**Figure 1:**
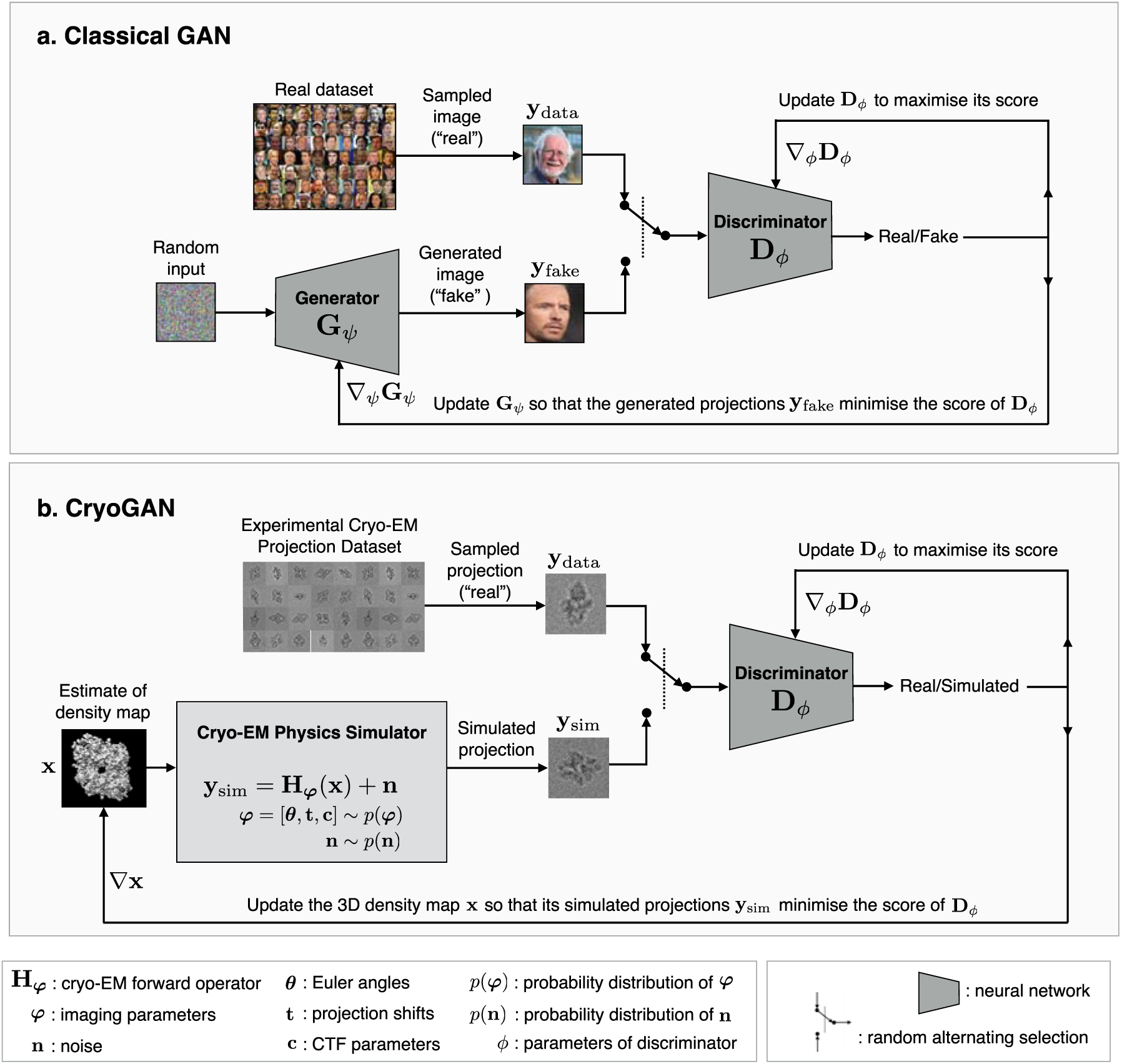
A schematic comparison between **(a)** a classical GAN architecture and **(b)** the CryoGAN architecture. Both frameworks rely on a deep adversarial learning scheme to capture the distribution of the real data. CryoGAN exploits this ability to look for the volume whose simulated projections have a distribution that matches the real data distribution. This is achieved by adding a “cryo-EM physics simulator” that produces measurements following a mathematical model of the cryo-EM imaging procedure. Importantly, CryoGAN does not rely on a first low-resolution volume estimate, but is initialized with a zero-valued volume. Note that, for both architectures, the updates involve backpropagating through the neural networks; those actions are not indicated here for the sake of clarity.

The CryoGAN architecture represents a complete change of paradigm for single-particle cryo-EM reconstruction. No estimation of the poses is attempted during the learning procedure; rather, the reconstruction is obtained through distributional matching performed in a likelihood-free manner. Hence, CryoGAN sidesteps many of the computational drawbacks associated with likelihood-based methods.

In practice, CryoGAN requires no prior knowledge of the 3D structure; its learning process is purely unsupervised. The user needs only to feed the particle images and estimates of the contrast transfer function (CTF) parameters. No initial volume is needed: the algorithm starts with a volume initialized with zeros. The CryoGAN framework is backed up by a comprehensive mathematical framework that provides guarantees on the recovery of the correct structure under a given set of assumptions that are often met in practice, at least to some degree of approximation.

We first assessed the performance and stability of CryoGAN on a synthetic *β*-galactosidase dataset, where we generated noisy projections *in silico*. The results demonstrate that our unsupervised reconstruction paradigm permits the recovery of a 8.6 Å structure (Figure 2). We then deployed CryoGAN on a real experimental *β*-galactosidase dataset (EMPIAR-10061) [17], reaching a resolution of 12.1 Å in 150 minutes in far more challenging conditions (Figure 3). These preliminary results provide a strong indication of the suitability of the CryoGAN framework for the reconstruction of real structures. On the implementation side, we expect to be able to improve the resolution of the reconstructions by taking advantage of the many recent technical developments and advances in the area of GANs. In the meantime, the preliminary results obtained with CryoGAN are encouraging and demonstrate the potential of adversarial-learning schemes in image reconstruction. The proposed paradigm opens many new perspectives in single-particle cryo-EM and paves the way for more applications beyond the present one.

**Figure 2:**
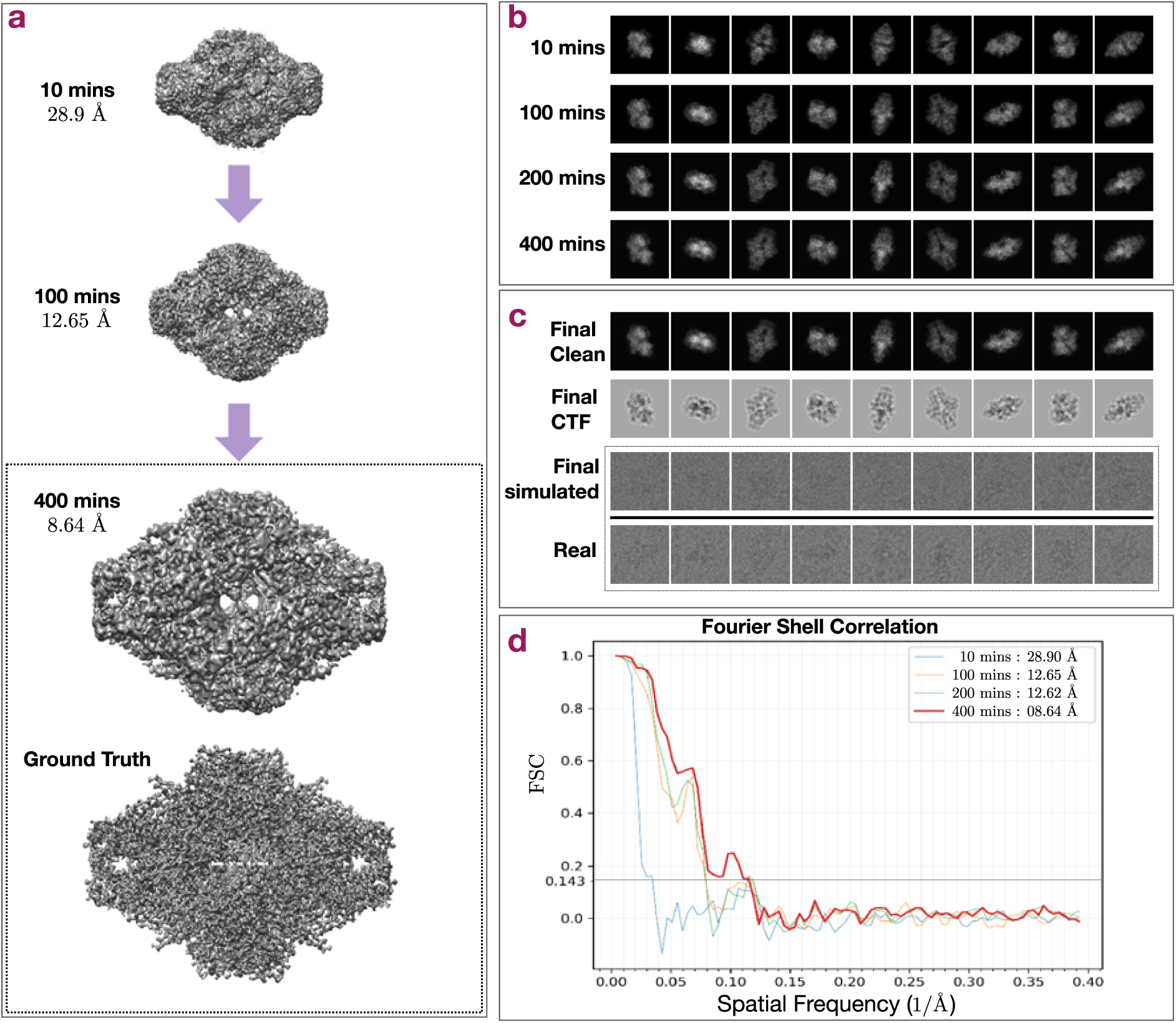
CryoGAN is applied on a synthetic projection dataset generated from a 2.5Å *β*-galactosidase volume. We refer to these synthetic projections as “real,” in contrast to the projections coming from CryoGAN, which we term “simulated.” **(a)** The volume is initialized with zeros and is progressively updated to produce projections whose distribution matches that of the real projections. **(b)** Evolution during training of some clean projections (*i*.*e*., before CTF and noise) generated by the cryo-EM physics simulator. **(c)** *Row 1* : Clean, simulated projections (before CTF and noise) generated at the final stage of training. *Row 2* : CTF-modulated simulated projections (before noise) generated at the final stage of training. *Row 4* : Real projections, for comparison. **(d)** FSC curves between the two reconstructed half-maps at different points during training.

**Figure 3:**
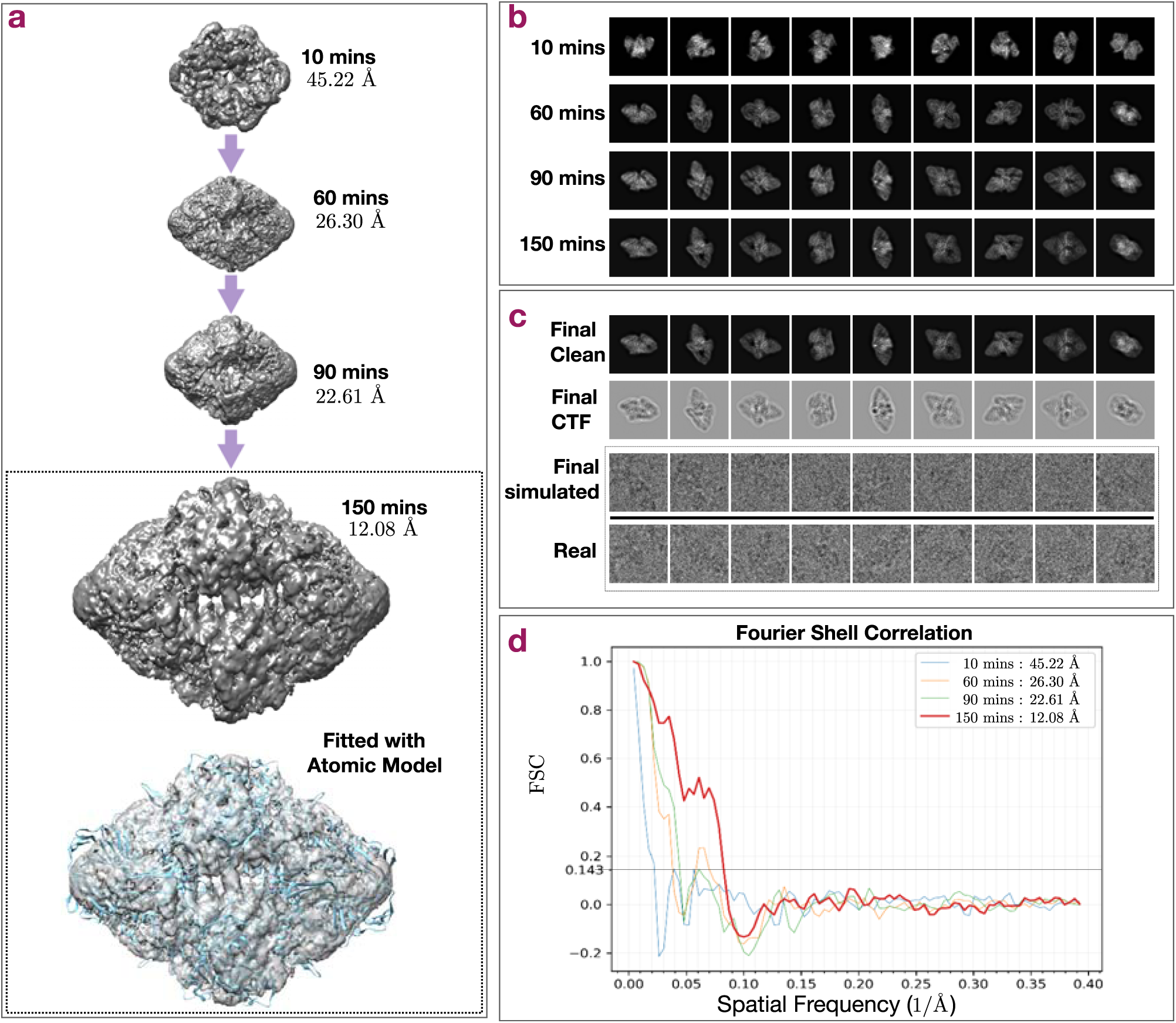
Evolution of CryoGAN while reconstructing the experimental *β*-galactosidase dataset (EMPIAR-10061) from [17]. **(a)** The volume is initialized with zeros and is progressively updated to produce projections whose distribution matches that of the experimental dataset. **(b)** Evolution during the training of the clean projections (*i*.*e*., before CTF and noise) generated by the cryo-EM physics simulator. **(c)** *Row 1* : Clean, simulated projections generated at the final stage of training. *Row 2* : CTF-modulated, simulated projections (before noise) generated at the final stage of training. *Row 3* : Simulated projections (with CTF and noise) generated at the final stage of training. *Row 4* : Real projections, for comparison. **(d)** FSC curves between the two reconstructed half-maps at different points during training.

## RESULTS

### The CryoGAN Algorithm

CryoGAN is like a standard GAN, except that the generator network is replaced by a cryo-EM physics simulator (Figure 1b). This simulator implements a mathematical model of the imaging procedure to produce a simulated measurement based on 1) the current volume estimate and 2) a random projection orientation. This image-formation model considers that the cryo-EM 2D measurement is the projection of the volume at that orientation, modulated by microscopy-related effects and corrupted by substantial additive noise.

The cryo-EM physics simulator is paired with a discriminator network whose architecture is similar to those used in standard GANs. The role of this discriminator in CryoGAN is to encourage the simulator to learn the 3D volume **x**_rec_ whose simulated dataset distribution *p*(**y**|**x**_rec_) matches that of the real dataset, *p*_data_(**y**). Mathematically, this equates to

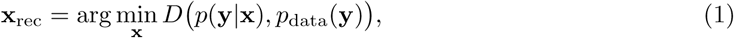

where *D* is an appropriate measure of distance between distributions, which the discriminator allows us to compute; in our implementation, *D* is the Wasserstein distance [18]. This formulation is based on a sound mathematical framework that provides guarantees on the recovery of the volume. In Theorem 1 (*Supplementary Note: Theoretical Guarantee of Recovery*), we show that, under certain constraints, the reconstructed volume is the same as the true volume up to rotation and reflection.

To perform the minimization (1), the CryoGAN algorithm alternates between updates of the discriminator and of the volume with stochastic gradient descent (SGD). The complete mathematical and algorithmic description of CryoGAN is given in *Online Methods*. The framework was implemented in Python using the PyTorch [19] package.

## Results on Synthetic Data

We first assessed the viability and performance of CryoGAN on a synthetic dataset that consists of 41,000 *β*-galactosidase projections, designed to mimic the EMPIAR-10061 data [17] in terms of noise level and CTF parameters. To create this dataset, we generated a 2.5 Å-resolution density map from the PDB entry (*5a1a*) of the protein and applied the forward model described in *Online Methods* to obtain projections modulated by CTF effects and corrupted by noise. We then randomly divided this dataset in two and applied the CryoGAN algorithm separately on both halves to generate half-maps. In the context of this experiment, we refer to these synthetic projections as “real,” in contrast to the projections coming from CryoGAN, which we term “simulated.” The details of the experimental setup are given in *Supplementary Note: Details of Experiments*.

We ran the CryoGAN algorithm for 400 minutes on an NVIDIA V100 GPU and obtained a reconstruction with a resolution of 8.64 Å (Figure 2a). Starting from a zero-valued volume, CryoGAN progressively updates the 3D structure so that its simulated projections (Figure 2b) reach a distribution that matches that of the real projections. These gradual updates are at the core of the deep adversarial learning scheme of CryoGAN. At each iteration of the algorithm, the gradients from the discriminator carry information about the current difference between the real projections and the simulated projections. These gradients are used by the cryo-EM physics simulator to update the volume so as to improve the fidelity of the simulated projections. Hence, at the end of its run, the volume learned by CryoGAN has simulated projections (Figure 2.c, Rows 1-3) that are similar to the real projections (Figure 2.c, Row 4) in a *distributional* sense. The evolution of the Fourier-shell correlation (FSC) between the reconstructed half-maps (Figure 2.d) testifies to the progressive increase in resolution that derives from this adversarial learning scheme.

We also evaluated the performance of CryoGAN on two additional synthetic datasets: one with a lower noise level and one with off-centered projections. Results were similar to those in Figure 2; further details are given in *Supplementary Discussion: Additional Results on Synthetic Data*.

### Results on Experimental Data (EMPIAR-10061)

We then deployed CryoGAN on 41,123 *β*-galactosidase projections (obtained from EMPIAR-10061 [17]) to assess its capacity to reconstruct real, experimental data. Here as well, we randomly divided the dataset in two and applied CryoGAN separately on both halves. The details of this experimental setup are given in *Supplementary Note: Details of Experiments*.

We ran CryoGAN for 150 minutes to obtain a 12.1 Å-resolution reconstruction using an NVIDIA V100 GPU. The results are displayed in Figure 3. The flexible architecture of CryoGAN permits the straightforward injection of prior knowledge on this specific imaging procedure into the reconstruction pipeline (*e*.*g*., the assumption of uniform pose distribution). Using this prior knowledge and its adversarial learning scheme, CryoGAN converges toward the reconstruction that best explains the statistics of the dataset (Figure 3a). As with the synthetic experiments, this is achieved by exploiting the gradients of the discriminator to update the simulator and the current volume estimate, so that, at later iterations, the simulated projections (Figure 3b) follow a distribution that better approaches that of the real dataset. Higher-resolution details are thus progressively introduced in the estimated volume throughout the run, as illustrated by the evolution of the FSC curves between the reconstructed half-maps (Figure 3d). This resulted in a 12.08 Å *β*-galactosidase structure whose simulated projections closely resemble the real ones (Figure 3c).

## DISCUSSION

We demonstrated the ability of CryoGAN to autonomously reconstruct 3D density maps through a purely data-driven adversarial learning scheme, which represents a complete change of paradigm for single-particle cryo-EM reconstruction. Capitalizing on the ability of deep-learning models to capture data distributions, the CryoGAN algorithm looks for the reconstruction that is most consistent with the measurements in a *distributional* sense. Hence, it is able to avoid the angular-assignment procedure altogether by directly exploiting the statistics of the provided dataset. CryoGAN is a purely unsupervised algorithm that requires minimal prior information and user input. It is backed up by a sound mathematical framework that gives guarantees on the recovery of the volume, provided that the image-formation model is valid (see *Online Methods* and *Supplementary Note: Theoretical Guarantee of Recovery*).

An important point is that CryoGAN is likelihood-free, which is in contrast to the iterative-refinement approaches used in most software packages [6, 9, 11]. This permits CryoGAN to bypass marginalization over the angles, a task that is inherent in these likelihood-based approaches, but that is cumbersome because it requires some form of angular assignment and/or the approximation of integrals by sums. CryoGAN also sidesteps many inconvenient processing steps, such as 2D alignment or classification, which further reduces the need for user-dependent inputs.

Our synthetic experiments demonstrate the ability of CryoGAN to resolve a structure so that its simulated projection distribution approaches that of the experimental particles. These results validate the CryoGAN paradigm and the viability of its current implementation. Indeed, without any prior training and starting from a zero-valued volume, the algorithm is able to autonomously capture the relevant statistical information from the dataset of noise-corrupted, CTF-modulated projections, and to progressively learn the volume that best explains these statistics. The results on the real *β*-galactosidase dataset further demonstrate the capacity of CryoGAN to perform reconstruction in challenging real imaging conditions.

### Roadmap for Future Work

Our implementation of CryoGAN is at the proof-of-concept stage and is bound to further improve. Beyond simple engineering tweaks (*e*.*g*., tuning the number of layers in the discriminator, testing different optimization strategies, or using Fourier methods to accelerate projection), we expect that several interesting developmental steps lie ahead.

A promising direction of research is the use of a coarse-to-fine strategy to reconstruct the volume progressively as the resolution improves. The motivation is that increased robustness during the low-resolution regime tends to have a positive impact on the convergence of the higher-resolution steps. Several GAN architectures rely on such approaches, such as the progressive GANs [20] and the styleGANs [21]. The benefits of multi-scale reconstruction could be considerable for CryoGAN, given the extremely difficult imaging conditions that prevail in single-particle cryo-EM and that make the convergence of optimization algorithms to good solutions particularly challenging. The core idea here would be to have the discriminator learn to differentiate between real and simulated distributions at a low resolution first, and then at successively higher ones. The impact on CryoGAN could be as important as the one it had on GANs, which progressed in just a few years from generating blurry facial images [16] to simulated images indistinguishable from real facial images [20, 21]. More generally, the upcoming tools and extensions in GAN architectures could bring significant gain in resolution to the CryoGAN implementation.

The performance of the cryo-EM physics simulator should also improve hand-in-hand with our ability to precisely model the physics behind single-particle cryo-EM with computationally tractable entities. At the moment, CryoGAN assumes that the noise is additive in its image-formation model. One could alternatively consider a Poisson-noise-based forward model [22, 23]. This would however require backpropagating through a Poisson distribution, a nontrivial operation.

Another interesting extension of the simulator would be to directly simulate the patches of nonaligned micrographs/frames (rather than the individual projections) and match their distribution to that of the raw dataset. Doing so would allow CryoGAN to bypass additional preprocessing tasks, in particular particle picking.

Similar to likelihood-based methods, the CryoGAN algorithm requires the specification of the distribution of poses. In the case of CryoGAN, one could also parameterize this distribution and learn its parameters during the reconstruction procedure, along the lines of [24]. The same approach could be used to calibrate the distribution of the translations of the projections.

On the theoretical side, we currently have mathematical guarantees on the recovery of volumes for which the assumed distribution of poses (be it uniform or not) matches the distribution of the real data. We have prior mathematical indications that this can also be achieved when there is a certain mismatch between the assumed distribution of poses and the actual one, given that an appropriate GAN loss is used.

Like all reconstruction algorithms, CryoGAN can fail if the dataset contains too many corrupted particle images, *e*.*g*., those with broken structures or strong optical aberrations. Several solutions could be deployed to handle excessive outliers in the data distribution. One approach would be to include a step that automatically spots and discards corrupted data so that the discriminator never sees them. Recent deep-learning-based approaches able to track outliers in data could prove useful in this regard [25].

Finally, the flexible architecture of CryoGAN should also facilitate its extension to the handling of structures with a continuum of conformational states, a feat often perceived as the greatest technical challenge ahead for single-particle cryo-EM [26]. A route of study we are currently exploring is to parametrize the different 3D conformations as outputs of an appropriate neural network, similar to the strategy used in ambient-GANs [27]. Hence, CryoGAN would learn the set of conformations whose projections are the most consistent with the acquired dataset, in a distributional sense here as well. Importantly, this would not only permit the recovery of the different conformations, but also give a probability on the structure of interest being in any given state in its conformational landscape, which could provide important biological insights.

While the spatial resolution of the CryoGAN reconstructions from real data is not yet competitive with the state-of-the-art, the algorithm is already able to steadily perform the hardest part of the job, which is to obtain a reasonable structure by using nothing save the particle dataset and CTF estimations. We believe that the aforementioned developments will help to bring the CryoGAN algorithm to the stage where it becomes a relevant contributor for high-resolution reconstruction in single-particle cryo-EM. We have laid out a roadmap for future improvements that should get us to this stage, and may eventually help us reconstruct dynamic structures.

## Acknowledgements

The authors would like to warmly thank Dr. Ricardo Righetto (University of Basel), Dr. Ricardo Adaixo (University of Basel), Prof. Henning Stahlberg (University of Basel, EPFL), and Dr. Sergey Nazarov (EPFL) for insightful discussions on single-particle cryo-EM. They are also thankful to Shayan Aziznejad (EPFL) and Dr. Quentin Denoyelle (EPFL) for useful feedback on mathematical developments.

This research was supported by the European Research Council (ERC) under the European Union’s Horizon 2020 research and innovation programme, Grant Agreement No. 692726 Global-BioIm: Global integrative framework for computational bio-imaging.

## ONLINE METHODS

### Image-Formation Model in Single-Particle Cryo-EM

We detail here the cryo-EM image formation model used in our implementation of the CryoGAN physics simulator.

We use the standard model for the single-particle cryo-EM imaging procedure [28]

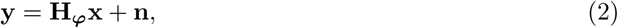

where **y** ∈ ℝ^*M*^ is a (vectorized) 2D projection; **x** ∈ ℝ^*V*^ is the (vectorized) 3D density; **H_*ϕ*_** ∈ ℝ^*M* ×*V*^ is the forward operator with parameters ***ϕ***; and **n** ∈ ℝ^*M*^ is an additive noise following a distribution *p*_**n**_. The imaging parameters ***ϕ*** comprise the projection (Euler) angles ***θ*** = (*θ*_1_, *θ*_2_, *θ*_3_), the projection shifts **t** = (*t*_1_, *t*_2_), and the CTF parameters **c** = (*d*_1_, *d*_2_, *α*_ast_), where *d*_1_ is the defocus-major, *d*_2_ is the defocus-minor, and *α*_ast_ is the angle of astigmatism.

The forward operator **H_*ϕ*_** is given by

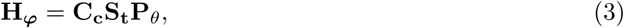

where **P_*θ*_** : ℝ^*V*^ → ℝ^*M*^ is a projection operator (mathematically speaking, the X-ray transform [29]), **S_t_** : ℝ^*M*^ → ℝ^*M*^ is a shift operator, and **C_c_** : ℝ^*M*^ → ℝ^*M*^ is a convolution operator. A more detailed description of the physics behind this image formation model **H_*ϕ*_** is given in *Supplementary Note: Image Formation*.

### Mathematical Framework of CryoGAN

The goal of single-particle cryo-EM reconstruction is to estimate a 3D density map **x**_rec_ whose projections are consistent with the observed projections (data) of the true density map **x**_true_.^1^

We write the conditional probability density function of a measurement **y** given a volume **x** by marginalizing over the imaging parameters ***ϕ***

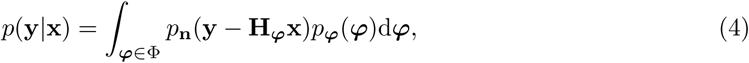

where *p*_***ϕ***_ is the distribution of the imaging parameters ***ϕ*** and Φ is the set of all their possible values. We denote a noiseless projection as **y**_noiseless_ = **H_*ϕ*_x**. In our formulation, the projections in the real dataset are samples of a distribution *p*_data_; hence, *p*(**y**|**x**_true_) = *p*_data_(**y**) assuming that the forward model is correct.

We demonstrate in Theorem 1 in *Supplementary Note: Theoretical Guarantee of Recovery* that two 3D volumes have identical conditional distributions if and only if they are identical, up to rotation and reflection. Hence, Theorem 1 implies that, for the reconstruction **x**_rec_ to satisfy **x**_rec_ = **x**_true_, it must also satisfy *p*(**y**|**x**_rec_) = *p*(**y**|**x**_true_). Thus, we can formulate the reconstruction task as the minimization problem

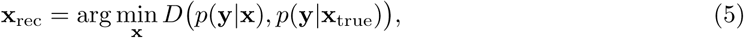

where *D* is some distance between two distributions. In essence, (5) states that the appropriate reconstruction is the 3D density map whose projection distribution is the most similar to the real dataset in a distributional sense. For the sake of conciseness, we shall henceforth use the notation *p*(**y**|**x**) = *p*_**x**_(**y**).

As distance in (5), we use the Wasserstein distance defined as

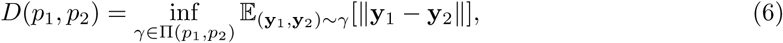

where Π(*p*_1_, *p*_2_) is the set of all the joint distributions *γ*(**y**_1_, **y**_2_) whose marginals are *p*_1_ and *p*_2_, respectively. Our choice is driven by works demonstrating that the Wasserstein distance is more amenable to minimization than other popular distances (*e*.*g*., total-variation or Kullback-Leibler divergence) for this kind of application [30]. Using (6), the minimization problem (5) expands as

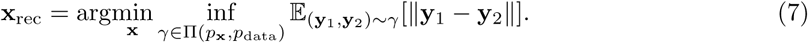

By using the formalism of [18, 30, 31], this minimization problem can also be stated in its dual form

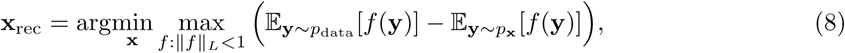

where ‖*f* ‖_*L*_ denotes the Lipschitz constant of the function *f* : ℝ^*M*^ → ℝ.

### Connection with Wasserstein GANs

Equation (8) falls under the framework of the generative adversarial networks (GANs) [16] called Wasserstein GANs (WGANs) [30]. In the standard WGAN representation, the function *f* is parameterized by a neural network **D**_*ϕ*_ with parameters *ϕ* that is called the discriminator. The task of this discriminator is to learn to differentiate between real samples (typically coming from an experimental dataset) and fake samples. The latter are produced by another neural network, called the generator, which aims at producing samples that are realistic enough to fool the discriminator. This adversarial-learning scheme progressively drives the WGAN to capture the distribution of the experimental data.

In CryoGAN, we exploit this capability of adversarial schemes to learn the volume **x** whose simulated projections follow the captured real-data distribution. To do so, we rely on a cryo-EM physics simulator, whose role is to produce projections of a volume estimate **x** using (2). These simulated projections then follow a distribution **y** ∼ *p*_**x**_. Hence, (8) translates into

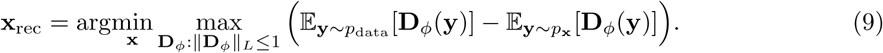

As proposed in [32], the Lipschitz constraint ‖**D**_*ϕ*_‖_*L*_ ≤ 1 can be enforced by penalizing the norm of the gradient of **D**_*ϕ*_ with respect to its input. This gives the final formulation of our reconstruction problem as

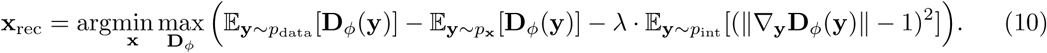

Here, *p*_int_ denotes the uniform distribution along the straight line between points sampled from *p*_data_ and *p*_**x**_, while *λ* ∈ ℝ_+_ is an appropriate penalty coefficient (see [32], Section 4).

### The CryoGAN Algorithm

Equation (10) is a min-max optimization problem. By replacing the expected values with their empirical counterparts (sums) [32], we reformulate it as the minimization of

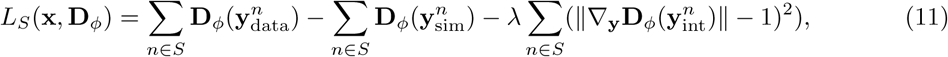

where *S* consists of either the full dataset *S*_full_ = {1, …, *N*_tot_} or a batch 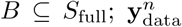 is a projection sampled from the acquired experimental dataset; 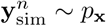 is a projection of the current estimate **x** generated by the cryo-EM physics simulator; and 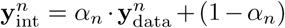, where *α*_*n*_ is sampled from a uniform distribution between 0 and 1.

In practice, we minimize (11) through stochastic gradient descents (SGD) using batches. We alternatively update the discriminator **D**_*ϕ*_ (for *n*_discr_ iterations) using an Adam optimizer [33] with gradient

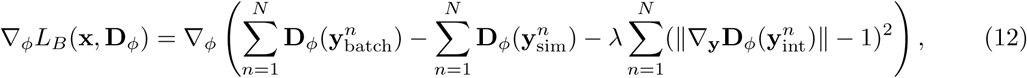

and the volume **x** (for 1 iteration) using the batch gradient,

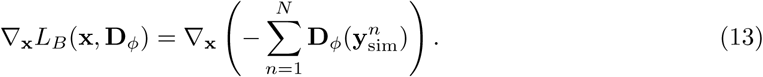

The pseudocode and a schematic view of the CryoGAN algorithm are given in Algorithm 1 and Figure 1b, respectively. We provide further details of the CryoGAN physics simulator and discriminator network in the next two sections.

#### Algorithm 1 Pseudocode for CryoGAN

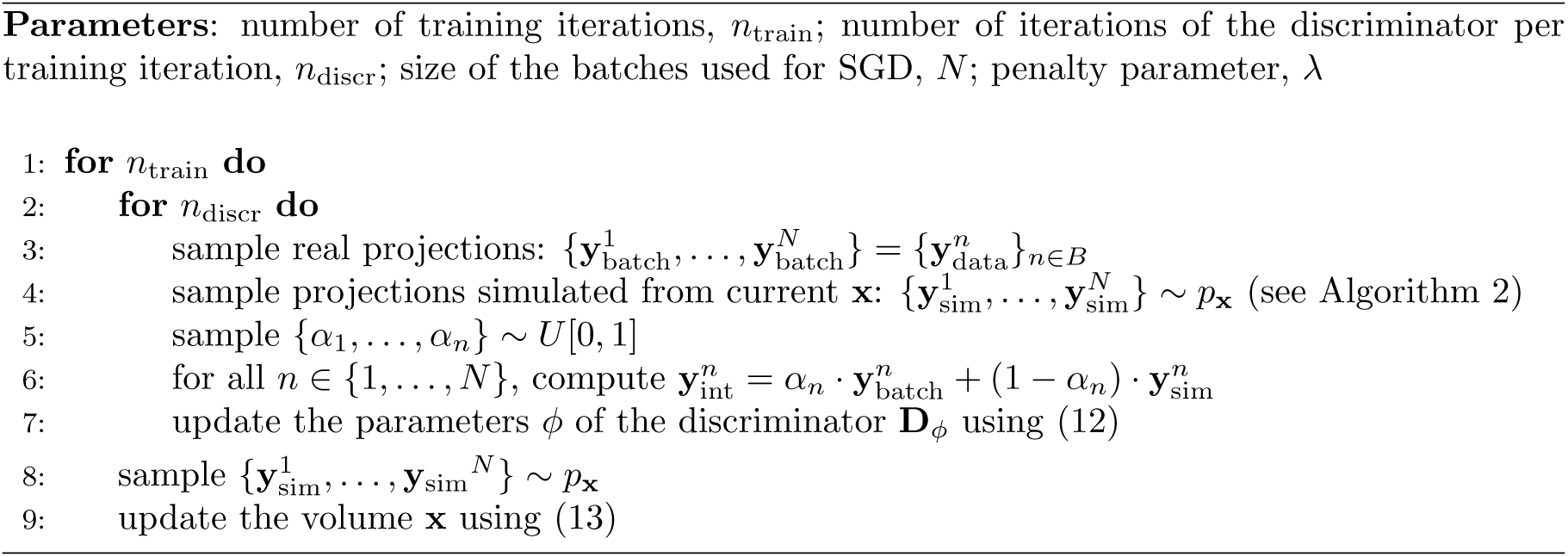

### The Cryo-EM Physics Simulator

The goal of the physics simulator is to sample **y**_sim_ ∼ *p*_**x**_(**y**); this is done in three steps. First, we sample the imaging parameters ***ϕ*** from the distribution *p*_***ϕ***_: ***ϕ*** ∼ *p*_***ϕ***_. Second, we generate noiseless CTF-modulated and shifted projections from the current volume estimate with **H_*ϕ*_**(**x**). Third, we sample the noise model to simulate noisy projections **y** = **H_*ϕ*_**(**x**)+ **n**, where **n** ∼ *p*_**n**_. We detail these steps in the following paragraphs, and a pseudocode of this cryo-EM physics simulator is given in Algorithm 2.

#### Algorithm 2 Pseudocode for Cryo-EM Physics Simulator

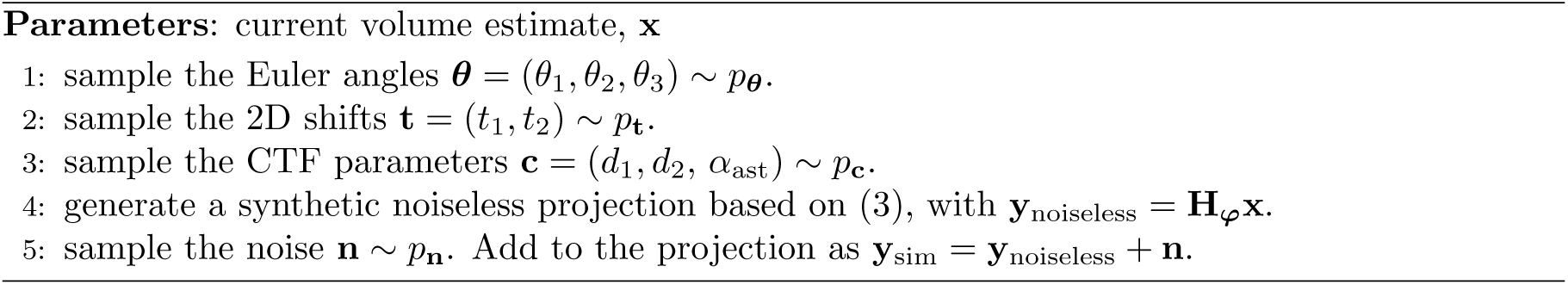

Recall that the set of imaging parameters is given by ***ϕ*** = (*θ*_1_, *θ*_2_, *θ*_3_, *t*_1_, *t*_2_, *d*_1_, *d*_2_, *α*_ast_). We first sample the Euler angles ***θ*** = (*θ*_1_, *θ*_2_, *θ*_3_) from a distribution *p*_***θ***_ decided *a priori* based on the acquired dataset. Similarly, the projection shifts **t** = (*t*_1_, *t*_2_) are sampled from the prior distribution *p*_**t**_. The CTF parameters **c** = (*d*_1_, *d*_2_, *α*_ast_) are sampled from the prior distribution *p*_**c**_. In practice, we exploit the fact that the CTF parameters can often be efficiently estimated for all micrographs. We then uniformly sample from the whole set of extracted CTF parameters.

We generate noiseless projections **y**_noiseless_ by applying **H_*ϕ*_** to the current volume estimate **x**. The projection operator **P_*θ*_** in (3) is implemented using the ASTRA toolbox [34].

The precise modeling of the noise is a particularly challenging feat in single-particle cryo-EM. To produce noise realizations that are as realistic as possible, we extract random background patches directly from the micrographs themselves, at locations where particles do not appear. For consistency, the noise patch added to a given noiseless projection is taken from the same micrograph that was used to estimate the CTF parameters previously applied to that specific projection. Additional details for this implementation are given in *Supplementary Note: Details of Experiments*.

### The CryoGAN Discriminator Network

The role of the discriminator is to differentiate between projections from the experimental dataset and projections generated by the cryo-EM physics simulator. The gradients from the discriminator (see (13) in Algorithm 1) carry information on the difference between the real and simulated projections at a given run-time. Those gradients are used by the simulator to update itself, thus improving on the realism of the simulated projections.

The discriminator network takes an image as input and outputs a scalar value. Its architecture is illustrated in Figure 4. It is composed of 8 layers: 6 convolutional blocks followed by 2 fully connected (FC) layers. Each convolutional block is made up of a convolutional layer followed by a max-pooling and a leaky ReLU (with negative slope of 0.1). The number of channels in each convolutional layer is 96, 192, 384, 768, 1536, and 3072, respectively. The filters in these layers are of size 3, and the padding size is 1. The max-pooling layer uses a kernel of size 2 with a stride of 2. This leads to a downsampling by a factor of 2. The output of the final convolutional block is then reshaped, fed into the FC layer with 10 neurons, and finally processed by a leaky ReLU. The resulting activations are fed to the last FC layer to output a scalar.

**Figure 4:**
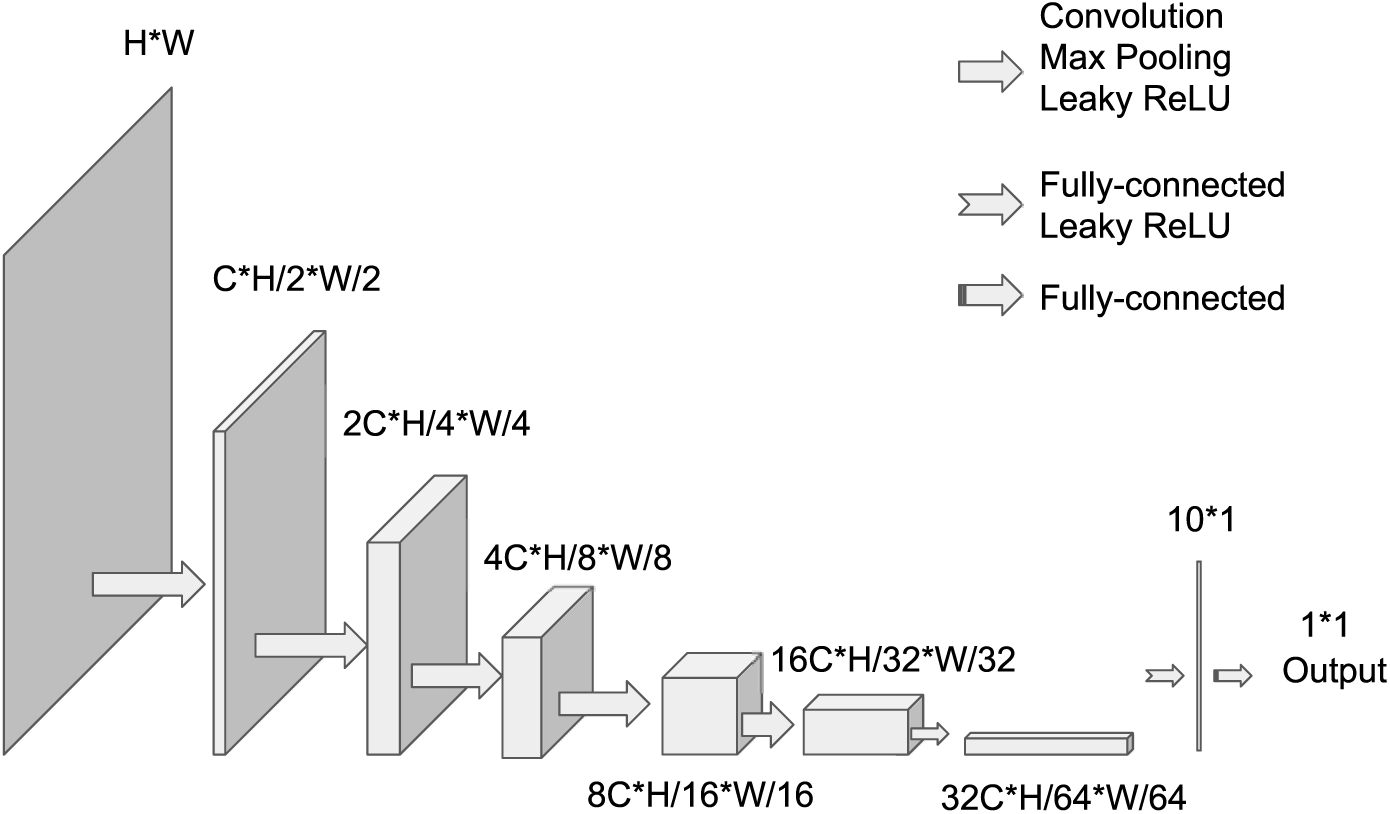
Architecture of the discriminator. The parameter for the channel size is *C* = 96 in every experiment. The input image with size *H* × *W* is successively processed and downsampled to output a scalar.

### Related Works

Related works fall into two main categories: current cryo-EM reconstruction methods and deep learning techniques that may apply to the cryo-EM pipeline; we now discuss each of these.

### Cryo-EM Reconstruction

The main challenge in cryo-EM reconstruction is that every particle has an unknown pose in its micrograph—if the poses were known, maximum-likelihood (ML) or maximum *a posteriori* (MAP) estimation of the volume could be performed by solving a standard linear inverse problem, where robustness would result from the large number of measurements which would counteract the low SNR of each measurement. One approach is to attempt to estimate the unknown poses iteratively. Pose estimation can be achieved with a variety of strategies, including the popular projection-matching approach [35, 36]. Whatever the method used, pose estimation is challenging because the SNR of individual projection images is extremely low. It also requires the estimation of additional parameters and the projection of the current reconstructed volume at a large number of poses and at every iteration of the reconstruction pipeline; ultimately, this is very computationally demanding.

Another approach is to formulate the reconstruction as a ML (or MAP) estimation problem in which the unknown poses are marginalized away [9, 11, 37]. This is attractive in that no extra parameter need to be estimated. The problem can then be solved using the expectation-maximization algorithm (e.g., [9,11]), where marginalization over poses during the so-called E-step is computationally expensive. Alternatively, the problem can be minimized using stochastic gradient descent (e.g., during the *ab initio* phase of [11]); here, the challenge is that the involved gradients require computations over all poses. For a more in-depth discussion, see [14, 15, 38]. For additional mathematical details on the relationship between likelihood-based methods and CryoGAN, see *Supplementary Note: CryoGAN vs. Likelihood-based Methods*.

Likelihood-free methods for cryo-EM reconstruction are relatively few. An early approach is [39], which proposes to reconstruct an *ab initio* structure such that the first few moments of the distribution of its theoretical cryo-EM measurements match the ones of the particles. However, the method assumes that the poses of the particles have a uniform distribution. This moment-matching technique has been recently extended in [24] to reconstruct an *ab initio* structure in the case of nonuniform pose distributions.

By contrast, our CryoGAN framework proposes to match the distribution of the theoretical cryo-EM measurements to that of the real projections, by which we mean all the moments and not just the first few. Moreover, our method works for any pose distribution of the particles provided it is known beforehand. Alternatively, one could rely on a parametric model of the pose distribution and use the backpropagation mechanism of neural networks to learn its parameters during the CryoGAN run; this technique is explored in [24].

### Deep Learning for Cryo-EM

Deep learning has already had a profound impact in a wide range of image-reconstruction applications [40–42]; however, their current utilization in cryo-EM is mostly restricted to preprocessing steps such as micrograph denoising [43] or particle picking [44–48]. A recent work uses neural networks to model continuous generative factors of structural heterogeneity [49]. However, the algorithm necessitates a pose-estimation procedure that relies on a merely conventional approach. Another recent work [25] uses a variational autoencoder trained using a discriminator-based objective to find a low-dimensional latent representation of the particles. These representations are then used to estimate the poses.

Deep learning is now extensively used to solve inverse problems in imaging [41, 50–52]. However, most methods are based on supervised learning and thus rely on training data. A GAN-based scheme that recovers the underlying distribution of the data from its noisy partial observations through a forward model was recently proposed in [27]. Finally, the reconstruction of a 3D structure (implicitly or explicitly) from its 2D viewpoints (and not projections) is an important problem in computer vision [53]. Many recent deep-learning algorithms have been used in this regard [54, 55]. While this problem is ostensibly similar to cryo-EM reconstruction, the measurement model for these problems is much less complicated than it is for cryo-EM and bares no relation to this modality.

## SUPPLEMENTARY MATERIALS

## Supplementary Discussion: Additional Results on Synthetic Data

The main synthetic experiment given in Figure 2 considers imaging conditions with a realistic noise level (−20 dB SNR). We performed additional experiments on synthetic *β*-galactosidase datasets to understand the effect of different imaging conditions on the quality of CryoGAN reconstruction. More precisely, we considered 1) the case with a lower noise level (−5.2 dB SNR) and 2) the case with a realistic noise level (−20 dB SNR) and translations (3% of the image size) in the projections. All the other conditions are identical to the main synthetic experiment. More details on these two additional experiments are provided in *Supplementary Note: Details of Experiments*.

The reconstructions obtained in the two cases reach a resolution of 7.53 Å and 10.8 Å, respectively, as shown in Supplementary Figure 1. As expected, the decrease of the noise level from -20 dB to -5.2 dB improves the reconstruction quality (from 8.64 Å to 7.53 Å). For the second case, the challenging presence of translations results in a slightly lower resolution. We believe that using a discriminator architecture that is invariant to shifts in the input image would improve the result in this case. Indeed, the discriminator would then be blind to such shifts in the real and simulated data, which would yield a reconstruction with a quality similar to the no-shift case.

**Supplementary Figure 1:**
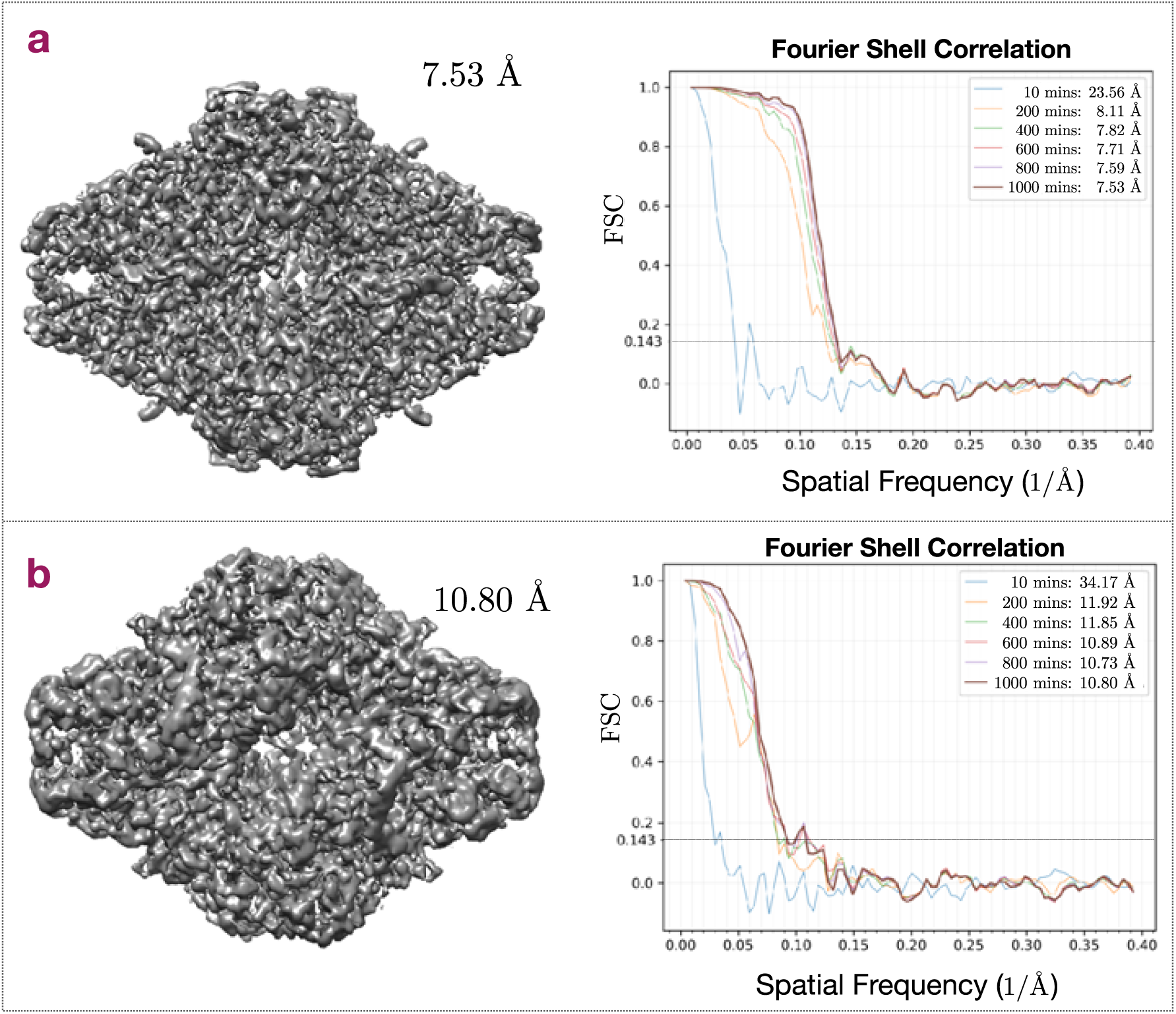
Additional CryoGAN reconstructions for synthetic datasets with different imaging conditions. **(a)** Reconstruction for a low noise case (−5.2 dB SNR). The corresponding evolution of the FSC curves with time is shown on the right. **(b)** Reconstruction for a realistic noise level (−20 dB SNR) and with translations (3% of the image size) in the data. The corresponding evolution of the FSC curves with time is shown on the right.

## Supplementary Note: CryoGAN vs. Likelihood-based Methods

As discussed in [15], most reconstruction approaches in SPA currently rely a on maximum-likelihood (ML) formulation [37] written as

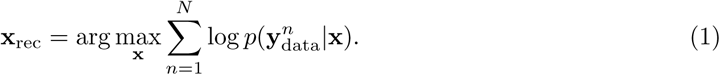

This formulation can be solved by using an expectation-maximization (ML-EM) algorithm or gradient descent. The former is preferred for iterative refinement [9], while a stochastic version of the latter is used for generating initial volume estimates in [11]. These techniques all involve the likelihood-estimation for each projection given the current volume estimate, which requires marginalizing the joint distribution *p*(**y**, *θ*|**x**) over the space of poses. In ML-EM, this is side-stepped in the expectation step (E-step) by computing the conditional distribution on the poses for each projection given the volume estimate. This conditional distribution is then used to update the volume in the marginalization step (M-step). Hence, all ML techniques require, either implicitly or explicitly, computations over a large number of poses for each projection.

For another perspective, ML techniques can be viewed as distribution-matching approaches. Specifically, (1) minimizes an empirical estimate of the Kullback-Leibler (KL) divergence between the real distribution *p*_data_(**y**) and the simulated distribution *p*(**y**|**x**), such that

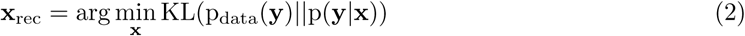

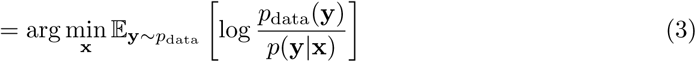

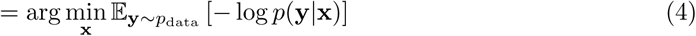

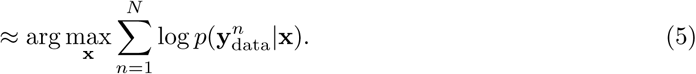

Hence, ML methods aim at finding the reconstruction whose simulated projection distribution matches that of the real data. In practice, this specific goal can be achieved by minimizing any suitable distance between these two distributions. By changing the distance, one can avoid the challenging likelihood computations that are inherent to the current ML methods, while preserving the theoretical guarantees that come with distribution matching (*e*.*g*., Theorem 1).

This is precisely the philosophy behind CryoGAN, which relies on the Wasserstein distance to formulate the distribution-matching task as

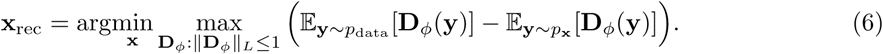

In this formulation, the two distributions indirectly interact through a common function: the discriminator network *D*_*ϕ*_. As a result, likelihood estimation is avoided; only a reliable sampler for each of the two distributions [56] is required. The samples from the real data distribution are readily available in the form of the acquired projection dataset, while the ones from the simulated distribution are generated by the cryo-EM physics simulator. Hence, CryoGAN is a likelihood-free technique with theoretical properties that are at least as good—if not better—than the ML ones. In fact, the Wasserstein distance is often easier to minimize than the KL divergence (*e*.*g*., due to the smoothness of the former) [30].

## Supplementary Note: Image Formation

For our forward model, we follow the developments in [28] (2.1-2.10), [57], and [23], which model the relationship between the 3D density map and a 2D measurement as linear.

The continuous-domain measurement can be given in the following form

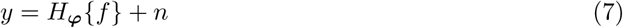

where *y* : ℝ^2^ → ℝ is the intensity measured on the image plane, *n* : ℝ^2^ → ℝ is the noise, *f* : ℝ^3^ → ℝ is the density map we aim to recover, and *H*_***ϕ***_ is the measurement operator dependent on the imaging parameters ***ϕ***. This model can be further decomposed as

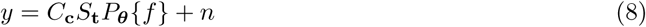

where ***ϕ*** = (***θ*, t, c**). We now discuss the three involved operators in more details.

### Projection Operator

The projection operation is given by [29]

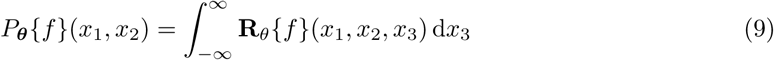

where **R_*θ*_** is the rotation matrix associated with ***θ*** and 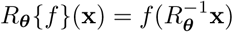.

### Shift Operator

The projection measurements are picked from the micrographs and can thus be off-centered. This is modelled via the shift operator which, for any *y*_*s*_ : ℝ^2^ → ℝ, yields

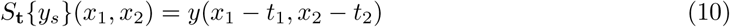

where **t** = (*t*_1_, *t*_2_).

### Convolution by CTF

The effect of the operator of *C*_**c**_ on any *y*_*c*_ : ℝ^2^ → ℝ is given in Fourier domain as

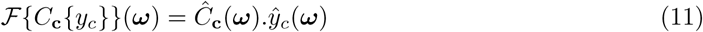

where *ℱ* is the Fourier transform and 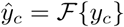. Its Fourier transform *Ĉ*_**c**_ (*i*.*e*., the CTF) is given by

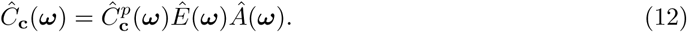

There, 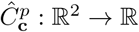 is the phase-contrast transfer function that takes the form

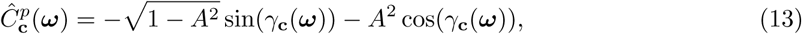

With

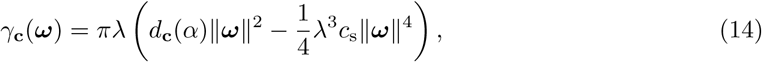

where *λ* is the electron wavelength, *c*_*s*_ is third-order spherical aberration constant, *α* is the phase of the vector ***ω***, and *d*_**c**_(*α*) is the defocus arising at the phase *α*. This defocus is given as

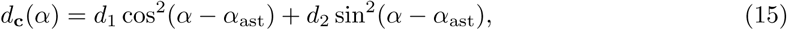

where *d*_1_ and *d*_2_ are the horizontal and vertical defocus, respectively, and *α*_ast_ is the reference angle that defines the azimuthal direction of axial astigmatism. The objective aperture function *Â* : ℝ^2^ → ℝ is given by

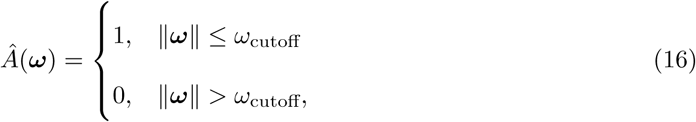

where 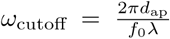 is the cutoff frequency, *f*_0_ is the focal length of the objective lens, and *d*_ap_ corresponds to the diameter of the aperture. The spatial and chromatic envelope function *Ê* : ℝ^2^ → ℝ is given by

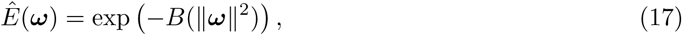

where *B* : ℝ^2^ → ℝ is a function influenced by chromatic aberration and spatial incoherence.

### Discretization

The discretization of *H*_***ϕ***_ results in **H_*ϕ*_**. This discretized measurement operator is itself decomposed of the discretized projection, shift, and convolution operation which are denoted by **P_*θ*_**, **S_t_**, and **C_c_**, respectively. The input to the operator **H_*ϕ*_** is a disretized version of the continous-domain 3D volume. This discretization of the 3D volume is done using a suitable basis function [58].

## Supplementary Note: Theoretical Guarantee of Recovery

The CryoGAN paradigm is supported by Theorem 1, which states that perfect recovery is possible from continuous-domain measurements. In the continuous domain, we have *y* = *H*_***ϕ***_*f* + *n* where *y* : ℝ^2^ → ℝ is the 2D measurement obtained from the 3D volume *f* and *n* is the noise. Here *H*_***ϕ***_ = *C*_**c**_*S*_**t**_*P*_***θ***_ is the continuous-domain forward operator, *P*_***θ***_ is the projection operator, *S*_**t**_ is the shift operator, and *C*_**c**_ is the operator for convolution with the CTF (see *Supplementary Note: Image Formation*).

**Theorem 1**. *Let y* = *H***_*ϕ*_***f* + *n as given in* (8) *with* ***ϕ*** = (***θ*, t, c**), *where* ***θ*** = (*θ*_1_, *θ*_2_, *θ*_3_) *are the projection angles*, **t** = (*t*_1_, *t*_2_) *are the shifts, and* **c** = (*d*_1_, *d*_2_, *α*_ast_) *are the CTF parameters (defocus-major, defocus-minor, and angle of astigmatism, respectively), f is the continuous-domain 3D volume, and y, n are continuous-domain 2D images. Let* ***θ*** ∼ *p*_***θ***_, **c** ∼ *p*_**c**_, **t** ∼ *p*_**t**_, *and* ℙ_*n*_ *be the probability measure associated with n. Moreover, assume that*

1. *the characteristic function* 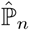 *of the noise probability measure* ℙ_*n*_ *is non-zero everywhere in its domain and n is pointwise defined everywhere in* ℝ^2^;
2. *the support of p*_**c**_ *is such that, for any* **c**_1_, **c**_2_ ∈ Support{*p*_**c**_} *and* **c**_1_ ≠ **c**_2_, *the fourier transform* 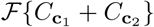 *is non-zero everywhere;*
3. *the volume f is nonnegative everywhere and has a bounded support; and*
4. *the probability distributions p*_***θ***_, *p*_**c**_, *and p*_**t**_ *are bounded*.

*Then, it holds that*

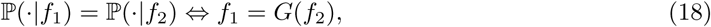

*where G is some member of the set of rotation-reflection operations*.

Before proving Theorem 1, we first comment on the assumptions. Assumption 1) is true for many noise distributions, including a white Gaussian noise filtered with any kernel of arbitrarily non-zero compact support. Assumption 2) is generally true as well. In fact, it is used to justify the application of Wiener filter to the clustered projections in classical cryo-EM reconstruction pipelines. Assumption 3) is true since the volume represents the density map, which is nonnegative. Also, the biological structures considered in cryo-EM have finite sizes.

*Proof of Theorem 1*. We denote the probability measure of *y*_noiseless_ = *H***_*ϕ*_***f* with P_noiseless_(*·*|*f*).

We shall prove the following in sequence:

1. ℙ (· |*f*_1_) = ℙ (· |*f*_2_) ⇔ ℙ_noiseless_(· |*f*_1_) = ℙ_noiseless_(· |*f*_2_),
2. ℙ_noiseless_(· |*f*_1_) = ℙ_noiseless_(· |**f**_2_) ⇔ *f*_2_ = *G*(*f*_1_).

For the first part we progress by noting that *y* = *y*_noiseless_ + *n*. Recall that the characteristic function of the probability measure associated to the sum of two random variables is the product of their characteristic functions. Mathematically,

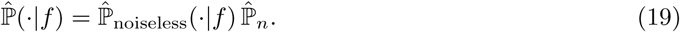

By Assumption (1, Theorem 1), we can now write that

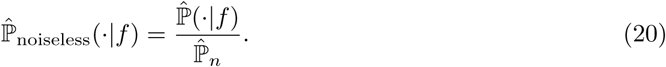

From (20), it is easy to see that ℙ (· |*f*_1_) = ℙ (· |*f*_2_) ⇔ ℙ_noiseless_(· |*f*_1_) = ℙ_noiseless_(· |*f*_2_). This concludes the first part.

For the second part, we now invoke the result from Theorem 4. It states that if *f*_2_ ≠ *G*(*f*_1_) for any *G* in the set of rotation and reflection operation, then the corresponding ℙ_noiseless_(· |*f*_1_) and ℙ_noiseless_(· |*f*_2_) are mutually singular. This means that the support of their intersection has zero measure. Since, we have ℙ_noiseless_(· |*f*_1_) = ℙ_noiseless_(· |*f*_2_) this means they are not mutually singular. This implies that *f*_2_ = *G*(*f*_1_) for some *G* in the set of rotation and reflection operation. This concludes the proof.

### Discrete-domain Extension

In practice, the cryo-EM measurements are acquired on a detector grid and are therefore discrete. CryoGAN reconstructs a voxel-domain **x**_rec_ such that *p*(**y**|**x**_rec_) = *p*(**y**|**x**_GT_), meaning that the corresponding probability measures are equal as well. Theorem 1 then holds approximately, up to some error that results from discretization of the measurements, forward model, and 3D density map. We leave a more thorough analysis of this error for future work.

In the following two sections, we build up to Theorem 4. First by analyzing the problem in the absence of the CTF and noise, and then by adding the CTF.

### Recovery in the Absence of Noise and CTF

In the absence of CTF and shifts the recoverability of *f* : ℝ^3^ → ℝ from its 2D projections obtained at unknown random poses is guaranteed by [59, Theorem 3.1]. We first go through the notations described in [59] before we state the required foundational result. We then extend [59, Theorem 3.1] to the case when the CTF and shifts are present.

### Notations and Preliminaries

Let *SO*(3) be the space of the special orthonormal matrices and 𝒟 be the Borel *σ*−algebra induced using the standard Riemannian metric on *SO*(3). Then, (*SO*(3), 𝒟) describes the measurable space of orthonormal matrices. Let 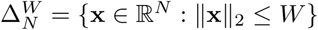 for some *W* ∈ ℝ^+^. By (*ℒ*_2_, *ℬ*), we denote the measurable space of all the square-integrable functions supported in 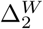 with Borel *σ*−algebra *ℬ* induced by the *L*_2_-norm. We denote by 𝔽 the set of all the functions supported in 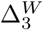, which are nonnegative and essentially bounded.

For any *f* ∈ 𝔽 and **A** ∈ *SO*(3), we denote 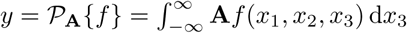 where **A***f* (**x**) = *f* (**A^−1^x**). Let *p*_**A**_ be a probability density on the space (*SO*(3), 𝒟). Note that there is a bijective mapping from ***θ*** in Theorem 1 and **A**. In fact, **A** represents the rotation matrix associated with the projection angle ***θ***.

We denote by Ψ the normalized Haar measure on (*SO*(3), 𝒟) and by Ψ_**A**_ the measure associated with *p*_**A**_ such that Ψ_**A**_[**·**] = (**a**∈**·**) *p*_**A**_(**a**)Ψ[d**a**].

For a given *f* ∈ 𝔽, the density *p*_**A**_ induces a probability measure ℙ_proj_(· |*f*) on the space (*ℒ*_2_, *ℬ*)through the mapping 𝒫_**A**_{*f*} such that

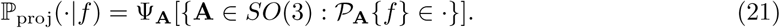

When *p*_**A**_ is uniform on *SO*(3), one has that

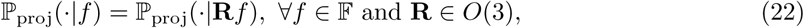

where *O*_3_ is the space of all orthogonal matrices such that det **A** ∈ {−1, 1}. The invariance in (22) is true since

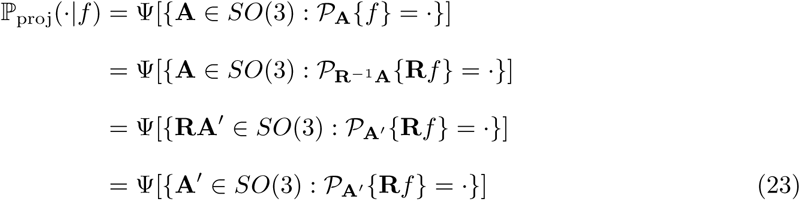

where **A**′ = **R**^−1^**A** and the last equality follows from the right invariance of Haar measure. We define *G*{𝔽} = {*γ*_*A*_ : *A* ∈ *O*_3_} such that

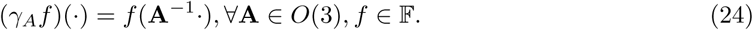

We define the shape [*f*] as an orbit of *f* under the influence of *G* such that [*f*] = {*γ*_**A**_*f* : *γ*_*A*_ ∈ *G*}. When *p*_**A**_ is uniform, the shape [*f*] is composed of all the rotations and reflections of *f*.

We can now restate [59, Theorem 3.1]. We discuss here the sketch of the proof given in [59].

**Theorem 2** ([59, Theorem 3.1]). *Let p*_**A**_ *be any bounded distribution on SO*(3) *and let the assumptions of Theorem 1 be true; then*, ∀*f, g* ∈ 𝔽,

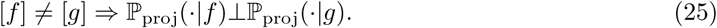

***Sketch of the Proof***. Without loss of generality, we provide the sketch of the proof for the case when *p*_**A**_ is uniform. For the case when *p*_**A**_ is nonuniform the argument remains the same provided that Ψ_**A**_ associated with the non-uniform distribution *p*_**A**_ is absolutely continuous with respect to Ψ (Ψ_**A**_ **≪**Ψ). This has been stated in [59]. Since we assume *p*_**A**_ to be bounded, this condition is satisfied. The only difference here with respect to the uniform distribution is that the orbit of *f* and *g* are more restricted than O(3).

The proof first uses in [60, Proposition 7.8] which we restate here as Proposition 3.

**Proposition 3** ([60, Proposition 7.8]). *Let f* ∈ 𝔽, *and let S*_**A**_ *be an uncountably infinite subset of SO*(3), *then f is determined by the collection* 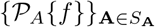 *ordered with respect to* **A** ∈ *S*_**A**_.

Note that this proposition assumes that the angle of the projections are known. Although in our case the angles are unknown, we shall see that this proposition will be useful.

We now want to determine how different ℙ_proj_(· |*f*) and ℙ_proj_(· |*g*) are for any given *f* and *g*. For this, we use the equality

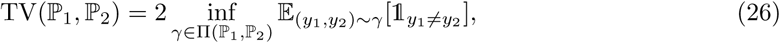

where TV is the total variation distance and Π(ℙ_1_, ℙ_2_) is the set of all the joint distributions *γ*(*y*_1_, *y*_2_) whose marginals are ℙ_1_ and ℙ_2_ [18]. In fact, 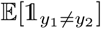 is equal to the probability of the event *y*_1_ ≠ *y*_2_ In our context, this translates into

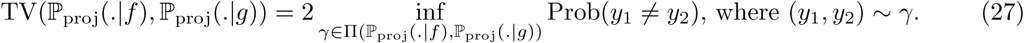

The optimum is achieved at the extremas which are sparse joint distributions and are such that the variable *y*_2_ is a function of *y*_1_. For any arbitrary joint distribution (or coupling) of this form, the proof then assigns a measurable function *h* : *SO*(3) → *SO*(3) such that (*y*_1_, *y*_2_) = (𝒫_**A**_{*f*}, 𝒫_*h*(**A**)_{*g*}) for **A** ∼ *p*_**A**_.

We can then write that

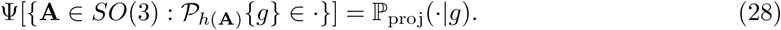

The task now is to estimate Prob(*y*_1_ ≠ *y*_2_), where (*y*_1_, *y*_2_) = (𝒫_**A**_{*f*}, 𝒫_*h*(**A**)_{*g*}) for **A** ∼ *p*_**A**_. *(Continuous h)*. When *h* is continuous, Proposition 3 implies that, if [*f*] ≠ [*g*], then

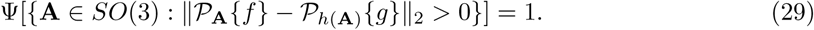

*(General h)*. When the function *h* is discontinuous, the proof uses Lusin’s theorem to approximate *h* by a continuous function. Lusin’s theorem states that, for any *δ >* 0, there exists an *h*_*δ*_ such that *h*(**A**) = *h*_*δ*_(**A**), ∀**A** ∈ *H*_*δ*_ and Ψ[*SO*(3)|*H*_*δ*_] < *δ*. This then leads to

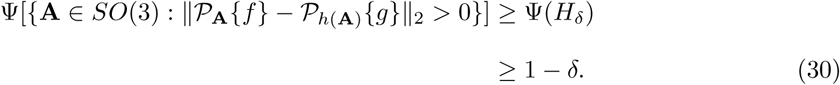

Since *δ* is arbitrarily small, the event {𝒫_**A**_{*f*} ≠ 𝒫_*h*(**A**)_{*g*}} has probability 1.

In conclusion, for any arbitrary coupling, the event {𝒫_**A**_{*f*} ≠ 𝒫_*h*(**A**)_{*g*}} has probability 1 if [*f*] ≠ [*g*]. This implies that, when [*f*] and [*g*] are not the same, the total-variation distance between 𝕡_proj_(· |*f*) and 𝕡_proj_(· |*g*) is 2. This ensures that the two probability measures are mutually singular meaning that the intersection of their support has zero measure. This concludes the proof.

### Recovery in the absence of Noise

We now extend the previous result to the case when the CTF is present. For the sake of simplicity we do not take into account shifts in the forward model. However, it is trivial to generalize the results to them since, shifts don’t change the information content in the projections but only their location.

We assume that **c** ∼ *p*_**c**_ such that the support of *p*_**c**_ is in some bounded region 𝒞 ⊂ ℝ^3^. We denote Ψ_**c**_[**·**] as the measure associated with *p*_**c**_ on the space 𝒞.

We denote by (*SO*(3) × 𝒞) the product space of *SO*(3) and 𝒞, while we denote by Ψ_**A**,**c**_ the measure on this product space. We then define

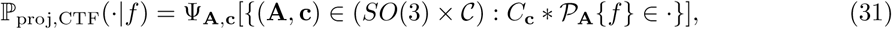

where *C*_**c**_ is the space-domain CTF given in (8).

**Theorem 4**. *Let p*_**A**_ *be a bounded probability distribution on SO*(3), *p*_**c**_ *be a distribution of the CTF with parameters* **c** ∈ 𝒞, *and let the assumptions of Theorem 1 be true; then*, ∀*f, g* ∈ 𝔽,

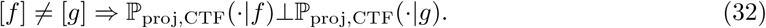

***Proof***. Similarly to the previous proof, we show that the TV distance between 𝕡_proj,CTF_(· |*f*) and 𝕡_proj,CTF_(· |*g*) is 2 when [*f*] and [*g*] are distinct. For simplification, we assume that *p*_**A**_ is uniform. (When this is not the case the proof essentially remains the same.) We need to show that Prob(*y*_1_ ≠ *y*_2_) = 1, where (*y*_1_, *y*_2_) ∼ *γ* for any arbitrary coupling *γ* of 𝕡_proj,CTF_(· |*f*) and 𝕡_proj,CTF_(· |*g*). For an arbitrary coupling *γ* such that Prob(*y*_1_ ≠*y*_2_) is minimum, we again assign *h* : (*SO*(3) × *C*) → (*SO*(3) × 𝒞) such that

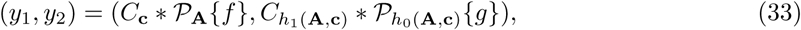

where **A** ∼ *p*_**A**_, **c** ∼ *p*_**c**_ and where *h*_0_ : (*SO*(3) × 𝒞) → *SO*(3) and *h*_1_ : (*SO*(3) × 𝒞) → 𝒞 are such that *h*(**A, c**) = (*h*_0_(**A, c**), *h*_1_(**A, c**)). This implies that

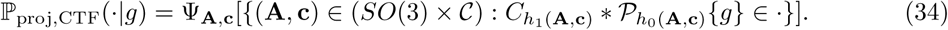

We now show that, for any *h*, the event {*y*_1_ ≠ *y*_2_} has probability 1. *(Continuous h)*. We first assume that *h* is continuous and use the same kind of technique as in the proof of [59, Theorem 3.1].

Since *SO*(3) is transitive, we can write that

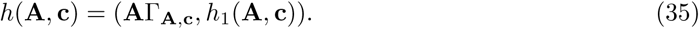

As *h* is continuous, so is Γ_**A**,**c**_. Let 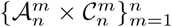 be a collection of *n* disjoint sets which creates the partition of (*SO*(3) × 𝒞). These partitions are such that for any *m*, there exists a *k*_*m*_ such that 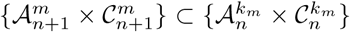. This means that, as *n* increases, the partitions become finer. We now define

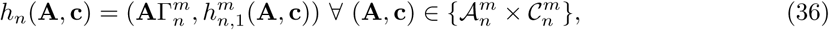

such that

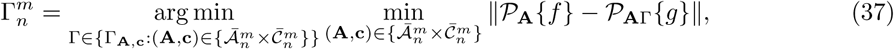

where 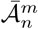 and 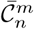 are the closures of 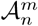 and 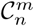, respectively. The sequence *h*_*n*_ converge to *h* as *n* → ∞. We denote

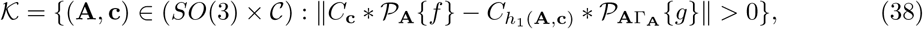

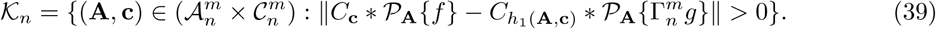

Similarly to [59, Theorem 3.1], we can then show that

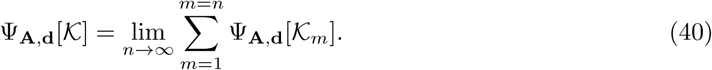

We invoke Proposition 5, which gives that 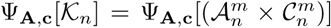. Therefore, Ψ_A,**d**_[𝒦] = Ψ_**A**,**c**_[(*SO*(3) × 𝒞)] = 1. This means that, when *h* is continuous, the event {*y*_1_ ≠ *y*_2_} has probability 1 if [*f*] ≠ [*g*].

*(General h)*. When *h* is discontinuous, we can invoke Lusin’s theorem to claim the same, similarly to Theorem 2. This means that, for any *h*, if [*f*] ≠ [*g*], then the probability of the event {*y*_1_ ≠ *y*_2_} is 1. Therefore, the TV distance between ℙ_proj,CTF_(· |*f*) and ℙ_proj,CTF_(· |*g*) is 2, yielding that ℙ_proj,CTF_(· |*f*)⊥ ℙ_proj,CTF_(· |*g*). This concludes the proof.

**Proposition 5**. *Let f, g* ∈ 𝔽, *A*^*I*^ ⊆ *SO*(3), 𝒞′ ⊆ 𝒞, Γ ∈ *SO*(3), *and*

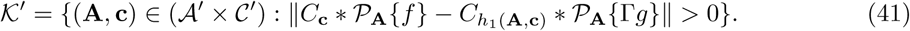

*Let the assumptions from Theorem 1 be true. Then, if* [*f*] ≠ [*g*], *it holds that*

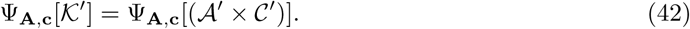

***Proof***. We show that Ψ_**A**,**c**_[𝒦^*’c*^] = 0, where (𝒦^*’c*^ ∪ 𝒦′) = (*A*′ × *C*′). We define the set 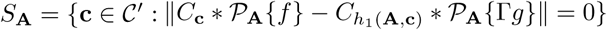. We define *S* _𝒜*”*_ = ∪ _𝒜∈ 𝒜*”*_ *S*_**A**_ for any 𝒜″ ⊆ 𝒜′. We define

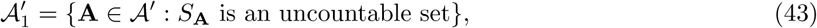

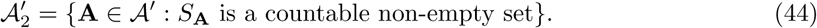

Note that 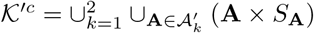. Then,

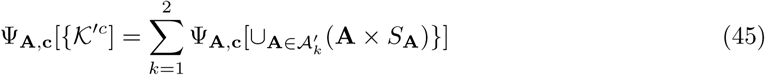

We now look at the two cases.

- *(When S*_**A**_ *is uncountable)*. For this case, we show that 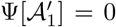. The main argument is that if this is not true, then it contradicts [*f*] ≠ [*g*].

For the sake of conciseness, we denote 𝒫_**A**_{*f*} by *I*_*f*_ and 𝒫_**A**_{Γ*g*} by *I*_*g*_. For any 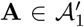, it holds that

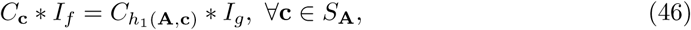

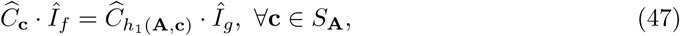

Where 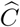, *Î*_*f*_, and *Î*_*g*_ are the Fourier transforms of **C**, *I*_*f*_, and *I*_*g*_, respectively.

We define 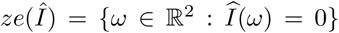, *ω*_*α*_ = {[(*r* cos *α, r* sin *α*)] : *r >* 0}, and *ze*_*α*_(*Î*) = *ze*(*Î*) ∩ *ω*_*α*_. From (47), we can write that

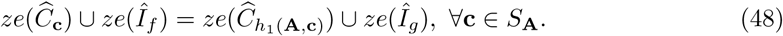

Two remarks are in order. Firstly, by assumption 2 of Theorem 1, 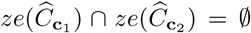 for **c**_1_ ≠ **c**_2_. (Remember that 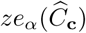 for any *α* ∈ [0, *π*] is nonempty (see “Image Formation Theory).) Secondly, by assumption 3 of Theorem 1, the supports of *f* and *g* are compact and nontrivial, so are the supports of *I*_*f*_ and *I*_*g*_. This means that their Fourier transforms Î_*f*_ and Î_*g*_ are analytic functions, which implies that there are infinitely many *α* such that the cardinality of the sets *ze*_*α*_(Î_*f*_) and *ze*_*α*_(Î_*f*_) is countable. We call the set of such *α* as *S*_*α*_. Now, we have that

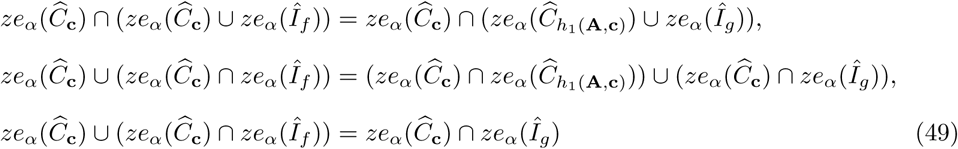

for all **c** ∈ *S*_**A**_ and *α* ∈ [0, *π*].

We can now write that

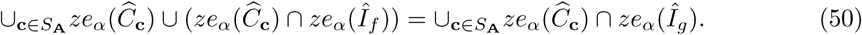

for any *α* ∈ *S*_*α*_. The set on the left hand side of (50) has an uncountably infinite cardinality since there are uncountably many **c** ∈ *S*_**A**_ and for each **c** there are distinct 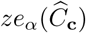. In return, the set in the right hand side of (50) is countable for a given *α* ∈ *S*_*α*_. Therefore, for any *α* ∈ *S*_*α*_, the two sets have different cardinality, which raises a contradiction. The only possible scenario in which (48) is true is when *h*_1_(**A, c**) = **c**. Using (47), we infer that 𝒫_**A**_{*f*} = 𝒫_**A**_{Γ*g*}. Therefore, for any 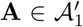, 𝒫_**A**_{*f*} = 𝒫_**A**_{Γ*g*}. However, 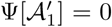 since, if this is not true, then [*f*] = [*g*] by Proposition 3.

Now note that

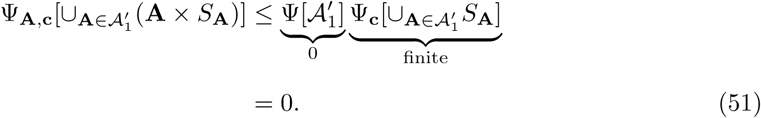

- *(When S*_**A**_ *is countable and nonempty)*. Since *S*_**A**_ is a countable set in this case, its elements have a bijection with natural numbers. We denote this bijection by *b* : 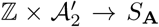. We denote by 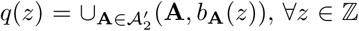. Note that *q*(*z*) is a graph of the function *b*(*z*, ·). Since it is a graph, Ψ_**A**,**c**_[*q*(*z*)] = 0.

We also have that 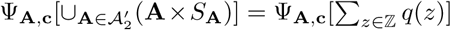. The latter vanishes since it is the measure of a countable addition of sets of measure zero. Hence, 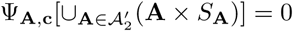.

This gives that 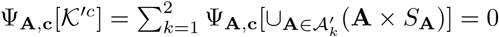, which concludes the proof.

## Supplementary Note: Details of Experiments

### Synthetic Data (Figure 2)

#### Dataset

We construct a synthetic cryo-EM dataset that mimics the experimental *β*-galactosidase dataset (EMPIAR-10061) from [17]. We generate 41,000 synthetic *β*-galactosidase projections of size 192 × 192 using our cryo-EM image-formation model (see *Online Methods*). For the ground-truth volume, we generate a 2.5 Å density map from PDB-*5a1a* atomic model using Chimera [61]. This gives a volume of size (302×233×163) with voxel size 0.637 Å. The volume is then padded, averaged, and downsampled to a size (180 × 180 × 180) with voxel size 1.274 Å. This corresponds to a Nyquist resolution of 2.548 Å for the reconstructed volume.

The projection poses are sampled from a uniform distribution over *SO*(3), where *SO*(3) is the group of 3D rotations around the origin of ℝ ^3^.

In order to apply random CTFs and noise, we randomly pick a micrograph in the EMPIAR-10061 dataset. We extract its CTF paramters using CTFFIND4 [62] and apply them to a clean projection. The parameter *B* of the envelope function of the CTF (see (17)) is chosen such that it decays to 0.2 at the Nyquist frequency. We then randomly select a background patch from the same micrograph to simulate noise. The noise is downsampled to size 192 × 192, and normalized to zero-mean, scaled and added to the projection. The scaling is such that the ratio of the energy of the signal to the energy of the noise (SNR) is kept at 0.01, which is equivalent to -20 dB. The dataset is then randomly divided into two halves. The algorithm is applied separately on both halves to generate the half-maps. The FSC between the two half maps is then reported using FOCUS [63, 64].

#### Generator Settings

We reconstruct a volume of size (184 × 184 × 184) voxels for each half dataset. The pixel size is 1.274 Å. The volumes are initialized with zeros. The D2 symmetry of *β*-galactosidase is enforced during reconstruction.

We use our image-formation model to simulate realistic projections from the current volume estimate at every CryoGAN iteration. The distribution of the imaging parameters is identical to the one used to generate the dataset. To add the noise on the CTF-modulated projections, we also keep the same approach as the one used to generate the dataset. However, we assume that the final SNR of each projection is unknown, leading us to learn the scaling parameter that controls the ratio between the projections and the noise patches.

We apply a binary spherical mask of size (184 × 184 × 184) on the learned volume. To handle the sharp transitions at the mask borders, we restrict the voxel values to lie above a certain value. This value changes as a function of position and iteration number: it increases linearly with the distance from the center of the volume to the border of the mask, from *V*_min_ to 0, and the value *V*_min_ changes as a function of iteration number, starting at 0 and decreasing to -2% of the current maximum voxel value. This promotes nonnegativity during the initial phases of reconstruction and increases the stability of the algorithm.

#### Discriminator Architecture

The architecture of the discriminator network is detailed in *Online Methods*. The discriminator is initialized identically for the two half datasets. All projections (*i*.*e*., the real projections and the ones generated by the simulator) are normalized to zero-mean and a unit standard-deviation before being fed to the discriminator.

#### General Settings

The adversarial learning scheme is implemented in Pytorch [19]. For the optimization, we use [33] (*β*_1_ = 0.5, *β*_2_ = 0.9, *E* = 10^−8^) with a learning rate of 10^−3^ and a batch size of 8. The algorithm is run for 40 epochs and the learning rate decreases by 1% at every epoch. The parameter for the gradient-penalty term (see (10)) is kept to *λ* = 0.001. The discriminator is trained 4 times for every training of the generator (*n*_discr_ = 4 in Algorithm 1).

For the back-propagations, the norm of the gradients for the discriminator are clipped to a maximal value of 10^8^. For the generator, the gradients for each pixel are clipped to a maximal value of 10^4^. The clipping values increase linearly from zero to those maxima in the first two epochs. Doing so improves on the stability of the adversarial learning scheme in the start, in particular, on that of the discriminator. All parameters are tuned for a fixed value range that follows from the normalization of all projections.

#### Computational Resources

The reconstruction is run on a Nvidia V100 GPU with 18GB memory. Each epoch lasts 10 minutes. The algorithm is run for 40 epochs which, in the current implementation, takes 400 minutes.

### Additional Synthetic Data (Supplementary Figure 1)

#### Synthetic dataset

The data is generated similarly to the main synthetic experiment, with the exception of some changes to the imaging conditions. In a first case, the noise level is set to a SNR of -5.2 dB. This corresponds to the energy of the noise being almost four times that of the signal. In the second case, the SNR is kept at -20 dB (the same as for the main experiment), but the projections are also translated. The translation (both horizontal and vertical) for each projection is sampled from a zero-mean symmetric triangular distribution whose total width is 6% of the image size from the centre. This corresponds to a shift of maximum 5 pixels from the centre in each direction.

#### Reconstruction Settings

The reconstruction settings for both cases are the same than the ones used in the main experiment, except for the few following differences. For the second case, the translations are also imposed, and the translation distribution is kept the same as the one used for generating the dataset. Furthermore, in both cases, the lower bound of the clipping value at the centre reaches -5% of the maximum voxel value of the volume. Finally, the algorithm is run for 100 epochs for both experiments.

### Experimental Data (Figure 3)

**Dataset**. The dataset consists of 41,123 *β*-galactosidase (EMPIAR-10061) projections extracted from 1,539 micrographs [17]. The projections of size (384 × 384) are downsampled to (192 × 192), with a pixel size of 1.274 Å. This corresponds to a Nyquist resolution of 2.548 Å for a reconstructed volume of size (180 × 180 × 180).

The dataset is randomly divided into two halves. The algorithm is applied separately on both halves to generate the half-maps. The defocus and astigmatism parameters of the CTF are estimated from each micrograph using Relion [9].

#### Generator Settings

We reconstruct a volume of size (180 × 180 × 180) voxels for each half dataset. The pixel size is 1.274 Å. The volumes are initialized with zeros. The D2 symmetry of *β*-galactosidase is enforced during reconstruction. A uniform distribution is assumed for the poses. The CTF parameters estimated with CTFFIND4 [62] are used in the forward model of the cryo-EM physics simulator. The parameter *B* of the envelope function of the CTF (see (17)) decays to 0.4 at the Nyquist frequency. The translations are set to zero.

To handle the noise, we randomly extract (prior to the learning procedure) 41,123 patches of size (384 × 384) from the background of the micrographs at locations where particles do not appear; this is done by identifying patches with the lowest variance. We extract as many noise patches per micrograph as we have particle images. Each noise patch is then downsampled to size (192 × 192) and normalized. Then, during run-time, the noise patches are sampled from this collection, scaled, and added to the simulated projections. For consistency, the noise patch added to a given simulated projection is taken from the same micrograph as the one that was used to estimate the CTF parameters previously applied to that specific projection. The scaling operation weights the contribution of the noise with respect to the projection signal. This is handled by multiplying the pixel values of the noise patches and the projections by two scalars that are learnt throughout the procedure. These two scalar values are the same for every pair of noise/projection images, so that the same amount of extracted noise is added to every simulated projection.

We apply a binary spherical mask of size (171 × 171 × 171) on the learned volume. To handle the sharp transitions at the mask borders, we enforce constraints on the masked volume. These are similar to those used in the synthetic experiment, with difference that the lower bound, *V*_min_ decreases to -5% of the maximum voxel value.

#### Discriminator Architecture

The architecture of the discriminator network is detailed in *Online Methods*. The discriminator is initialized identically for the two half datasets. The projection images (real and simulated) are smoothed with a Gaussian kernel before being fed to the discriminator. The standard deviation of the kernel is initially set at 2 and changes in every iteration so that it decreases by a total of 2% in each epoch.

#### General Settings

The adversarial learning scheme is implemented in Pytorch [19]. For the optimization, we use [33] (*β*_1_ = 0.5, *β*_2_ = 0.9, *E* = 10^−8^) with a learning rate of 10^−3^ and a batch size of 8. The algorithm is run for 15 epochs and the learning rate decreases by 8% at every epoch. The parameter for the gradient-penalty term is kept to *λ* = 1 (see (10)). The discriminator is trained 4 times for every training of the generator (*n*_discr_ = 4 in Algorithm 1).

For the back-propagations, the norm of the gradients for the discriminator are clipped to a maximal value of 10^7^. For the generator, the gradients for each pixel are clipped to a maximal value of 10^3^. The clipping values increase linearly from zero to those maxima in the first two epochs. Doing so improves on the stability of the adversarial learning scheme in the start, in particular, on that of the discriminator. The gradients that correspond to the learning of the scaling ratios between the noise and projection images are clipped to a value of 10.

#### Computational Resources

The reconstruction is run on a Nvidia V100 GPU with 18GB memory. Each epoch lasts 10 minutes. The algorithm is run for 15 epochs, which takes 150 minutes.

## Footnotes

1 We assume here conformational homogeneity of the underlying structure. The possible extension of CryoGAN to multiple conformations is discussed in *Discussion*.

